# RNA genome conservation and secondary structure in SARS-CoV-2 and SARS-related viruses

**DOI:** 10.1101/2020.03.27.012906

**Authors:** Ramya Rangan, Ivan N. Zheludev, Rhiju Das

## Abstract

As the COVID-19 outbreak spreads, there is a growing need for a compilation of conserved RNA genome regions in the SARS-CoV-2 virus along with their structural propensities to guide development of antivirals and diagnostics. Using sequence alignments spanning a range of betacoronaviruses, we rank genomic regions by RNA sequence conservation, identifying 79 regions of length at least 15 nucleotides as exactly conserved over SARS-related complete genome sequences available near the beginning of the COVID-19 outbreak. We then confirm the conservation of the majority of these genome regions across 739 SARS-CoV-2 sequences reported to date from the current COVID-19 outbreak, and we present a curated list of 30 ‘SARS-related-conserved’ regions. We find that known RNA structured elements curated as Rfam families and in prior literature are enriched in these conserved genome regions, and we predict additional conserved, stable secondary structures across the viral genome. We provide 106 ‘SARS-CoV-2-conserved-structured’ regions as potential targets for antivirals that bind to structured RNA. We further provide detailed secondary structure models for the 5’ UTR, frame-shifting element, and 3’ UTR. Last, we predict regions of the SARS-CoV-2 viral genome have low propensity for RNA secondary structure and are conserved within SARS-CoV-2 strains. These 59 ‘SARS-CoV-2-conserved-unstructured’ genomic regions may be most easily targeted in primer-based diagnostic and oligonucleotide-based therapeutic strategies.

## Introduction

Severe Acute Respiratory Syndrome Coronavirus 2 (SARS-CoV-2) has caused a rapidly expanding global pandemic, with the COVID-19 outbreak responsible at this time for over 600,000 cases and 25,000 deaths. The emergence of this pandemic has revealed an urgent need for diagnostic and antiviral strategies targeting SARS-CoV-2. Like other coronaviruses, SARS-CoV-2 is a positive sense RNA virus, with a large RNA genome approaching nearly 30 kilobases in length. Its RNA genome contains protein-coding open reading frames (ORFs) for the viral replication machinery, structural proteins, and accessory proteins. The genome additionally harbors various *cis*-acting RNA elements, with structures in the 5’ and 3’ untranslated region (UTRs) guiding viral replication, RNA synthesis and viral packaging.^1^ Conserved RNA elements offer compelling targets for diagnostics. In addition, such RNA elements may be useful targets for antivirals, a concept supported by the recent development of antisense oligonucleotide therapeutics and small-molecule RNA-targeting drugs for a variety of targets across infectious and chronic diseases.^2–4^

Conserved structured RNA regions have already been shown to play critical functional roles in the life cycles of coronaviruses. Most coronavirus 5’ UTR’s harbor at least four stem loops, with many showing heightened sequence conservation across betacoronaviruses, and various stems demonstrating functional roles in viral replication.^5^ Furthermore, RNA secondary structure in the 5’ UTR exposes a critical sequence motif, the transcriptional regulatory sequence (TRS), that forms long-range RNA interactions necessary for facilitating the discontinuous transcription characteristic to coronaviruses.^6^ Beyond the 5’ UTR, the frame-shifting element (FSE) in the first protein-coding ORF (ORF1 ab) includes a pseudoknot structure that is necessary for the production of ORF1 a and ORF1 b from two overlapping reading frames via programmed ribosomal −1 frame-shifting.^7^ In the 3’ UTR, mutually exclusive RNA structures including the 3’ UTR pseudoknot control various stages of the RNA synthesis pathway.^8^

Beyond these canonical structured regions, the RNA structure of the SARS-CoV-2 genome remains mostly unexplored. Unbiased discovery of other conserved regions and/or structured regions in the virus has the potential to uncover further functional *cis*-acting RNA elements. Here, we analyze RNA sequence conservation across SARS-related betacoronaviruses and currently available SARS-CoV-2 sequences, and we identify structured and unstructured regions that are conserved in each sequence set; these intervals can provide starting points for a variety of diagnostic and antiviral development strategies (Fig. 1). To identify structured regions, we predict maximum expected accuracy structures around conserved regions and report the support of these single structures from predictions of each RNA’s structural ensemble. We additionally identify thermodynamically stable secondary structures across the whole genome, finding that currently known structures fall within these predictions, but also identifying various new candidate structured regions. We pinpoint unstructured genome intervals by identifying bases with low average base-pairing probabilities. Finally, we present secondary structure models for key RNA structural elements of SARS-CoV-2 annotated in the betacoronavirus family.

**Figure 1:**
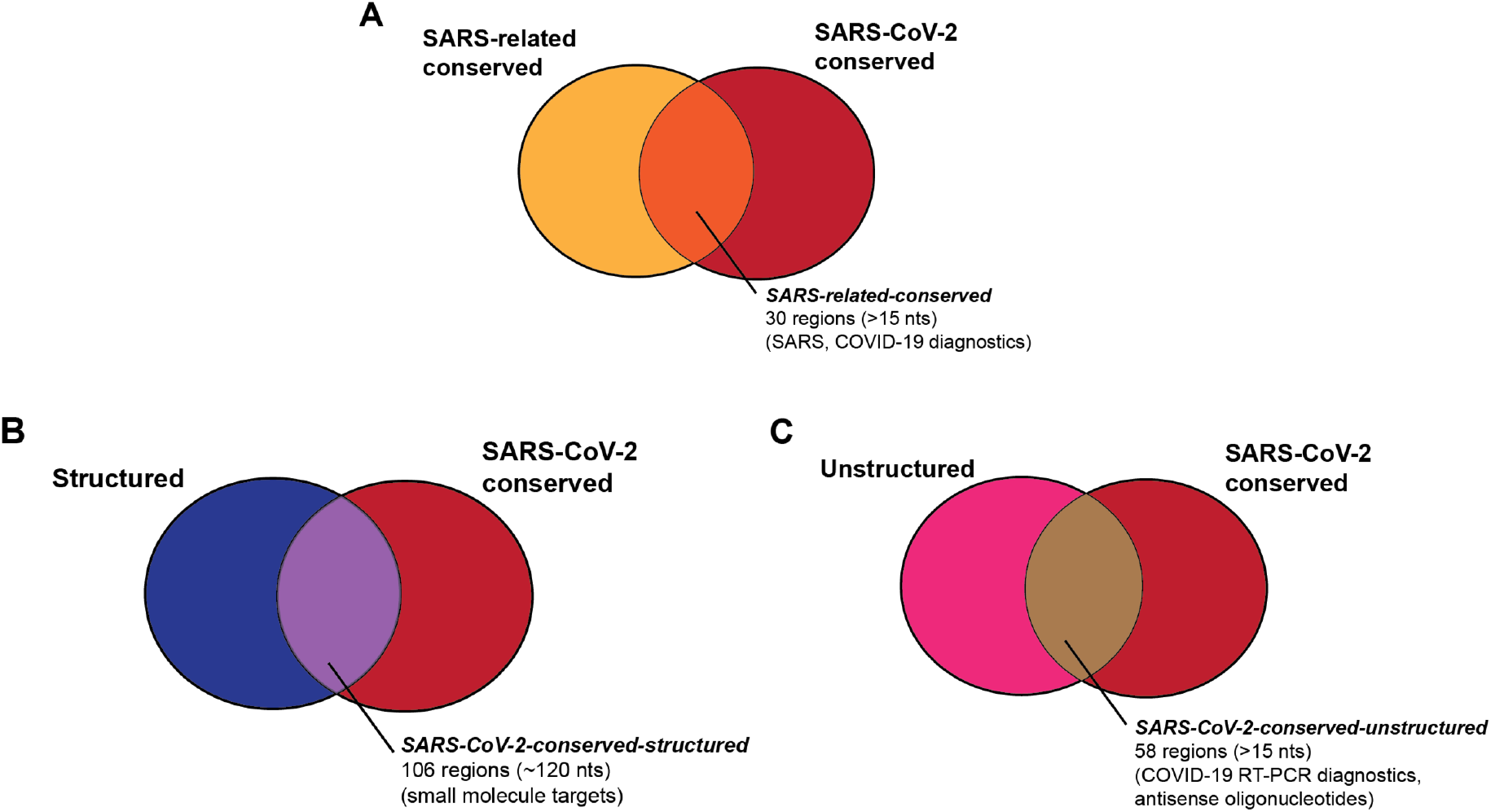
We aim to provide a series of genome regions in SARS-CoV-2 that are useful for a variety of diagnostic and therapeutic strategies, including regions that are (A) conserved in SARS-related betacoronaviruses and SARS-CoV-2 sequences (Table 1), (B) regions that are structured and conserved in SARS-CoV-2 sequences (Table 2), and (C) regions that are unstructured and conserved in SARS-CoV-2 sequences (Table 3).

## Results

### RNA sequence conservation in SARS-related betacoronaviruses and SARS-CoV-2

To identify potential regions of conserved RNA secondary structure in the virus, we located stretches of the SARS-CoV-2 genome with high RNA sequence conservation across SARS-related betacoronavirus full genome sequences. By identifying regions with high RNA sequence conservation as a first step, we reasoned that we would be more likely to filter for functionally relevant structures that must be conserved through virus evolution and thereby discover targets that are potentially less likely to develop resistance against therapeutics or to escape diagnosis as the virus evolves. To ensure reasonable numbers of sequences while still focusing on conservation and structure patterns most relevant to the current pandemic, we chose to analyze not all betacoronaviruses but a subgroup of SARS-related betacoronaviruses. These include SARS, SARS-CoV-2, and SARS-related bat coronaviruses, but not MERS, MHV, or other betacoronaviruses which have been classified into distinct subgroups based on different sequence and structure features in, for example, their 5’ UTR’s.^9^

We carried out this analysis beginning with three different sequence alignments. Each captures a range of complete genome sequences across the SARS-related betacoronviruses, but differ in the total number of sequences and in the redundancy of those sequences, as follows:

1. The first multiple sequence alignment (**SARSr-MSA-1**) was computed by aligning sequences curated by Ceraolo and Giorgi,^10^ filtered by including only the reference genome sequence NC_0405512.2^11^ from the SARS-CoV-2 sequence set, removing the two MERS sequences, and leaving in all remaining betacoronavirus whole genome sequences. This alignment captures a range of SARS-related bat coronavirus and SARS sequences with only 11 sequences. These sequences correspond well to the SARS-related group defined in Gorbalenya, Baker, *et al^12^*
2. The second MSA (**SARSr-MSA-2**) was obtained from BLAST by searching for the 100 complete genome sequences closest to the SARS-CoV-2 reference genome. This alignment captures a larger set of SARS-CoV-2, SARS, and bat coronavirus sequences than SARSr-MSA-1 but includes many sequences with high pairwise similarity.
3. The final MSA (**SARSr-MSA-3**) was obtained by locating all complete genome betacoronavirus sequences from the NCBI database, and removing mutually similar sequences with a 99% sequence conservation cutoff. With 180 sequences with at most 99% pairwise sequence similarity, this MSA captures a broader set of betacoronaviruses than SARSr-MSA-1 and SARSr-MSA-2 but is more challenging to align due to higher sequence diversity.

We computed conserved regions as contiguous stretches of 15 nucleotides or longer that were 100% conserved (cutoff for SARSr-MSA-1), 98% conserved (cutoff for SARSr-MSA-2), or 54% conserved (cutoff for SARSr-MSA-3). Searching for conserved regions of 15 nucleotides or more enables the design of antisense oligonucleotides that fall within these stretches. The sequence conservation cutoffs chosen ensured that at least 75 candidate conserved stretches were used for further structure analysis for each MSA. When calculating sequence conservation at the 5’ and 3’ sequence ends of the sequence, we did not include sequences that included only leading or trailing sequence deletions up to that point to avoid sequencing artefacts.

In Fig. 2, we depict conserved regions (100% conservation cutoff, SARSr-MSA-1) alongside the genome coordinates for the reference SARS-CoV-2 sequence. We observe intervals of conservation in the 5’ UTR and 3’ UTR genome regions, as expected based on prior work demonstrating sequence conservation surrounding structured RNA elements in these regions,^13^ but we also noted stretches of RNA sequence conservation within some viral ORFs.

**Figure 2:**
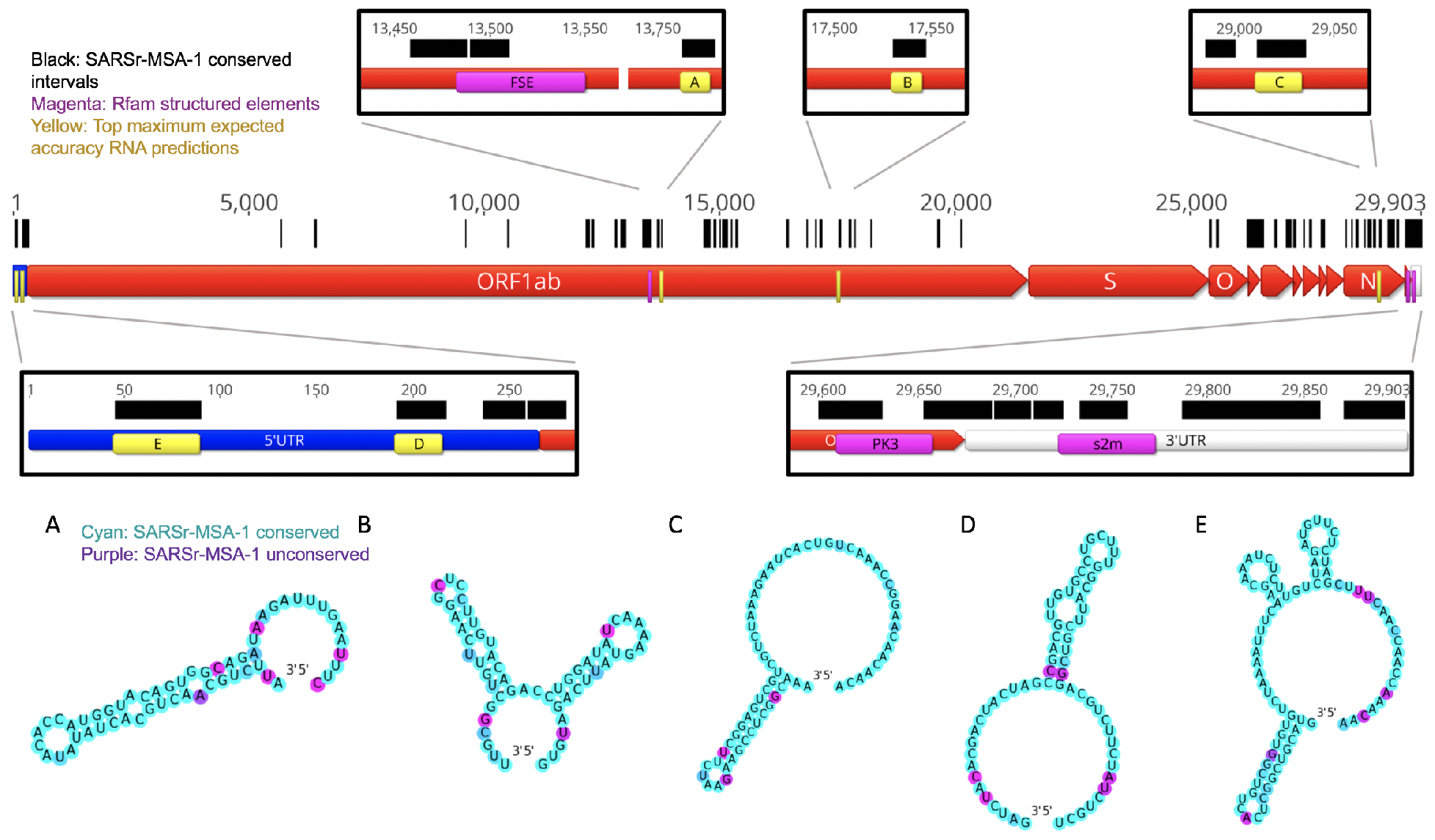
In black we annotate SARSr-MSA-1 conserved regions of the genome, superimposed on SARS-CoV-2 genome ORFs. We depict the top secondary structures as ranked by Matthews correlation coefficient that overlap with these conserved regions, ordered from A to E. Regions A to E are annotated on the genome in yellow and are located at genome positions: A:13743-13798, B:17511-17566, C:28990-29054, D:172-236, E:26-109. Secondary structures are colored by sequence conservation in SARSr-MSA-1 (cyan = more conserved, purple = less conserved). In magenta are depicted curated Rfam families present in coronaviruses, including the frame-shifting element (FSE), the 3’ UTR pseudoknot (PK3), and the 3’ stem-loop II-like motif (s2m). Figures prepared in Geneious^41^ and draw_rna (https://github.com/DasLab/draw_rna).

Interestingly, in SARSr-MSA-1 and SARSr-MSA-2 we found that conserved stretches overlapped with previously curated Rfam^14^ families for *Coronaviridae* RNA secondary structures: the frameshifting stimulation element (Rfam family RF00507), the 3’ UTR pseudoknot (Rfam family RF00165), and the 3’ stem-loop II-like motif (Rfam family RF00164) (Fig. 2). Locations for the frameshifting stimulation element, 3’ UTR pseudoknot, and 3’ stem-loop II-like motif were confirmed using Infernal,^15^ with all regions discovered at an E<10^-4^ threshold. We also found overlap between conserved stretches and additional 5’ UTR structures that have been established for previous coronaviruses, including the original SARS virus, including stem loops 2-3 (SL2-3) and stem loop 5 (SL5).^16^ These five known RNA structures overlap with conserved regions more than expected; in 10,000 random trials, the chance that five randomly chosen intervals of these lengths all overlap with the conserved regions from SARSr-MSA-1 or SARSr-MSA-2 is less than 0.0003. The enrichment of known RNA structures in these conserved regions suggests that other conserved regions may also harbor RNA structures.

To further tighten this list of conserved sequences to ones most relevant to the current COVID-19 outbreak, we analyzed whether sequence regions conserved across SARS and bat coronaviruses remain conserved in the SARS-Cov-2 strains, most of which emerged after our analysis above (Fig. 1A). We determined the conservation of conserved genome regions from SARSr-MSA-1 across SARS-CoV-2 sequences as of deposition date 03-18-20. For this analysis, we obtained two wholegenome multiple sequence alignments, keeping only full-length genome sequences of at least 29,000 nucleotides in both cases: the first includes 103 NCBI sequences (**SARS-CoV-2-MSA-1**), and the second includes 739 sequences deposited to GISAID^17^ (**SARS-CoV-2-MSA-2**). We noted conserved regions in the betacoronavirus alignment SARSr-MSA-1 were more likely to be at least 99% conserved in both SARS-CoV-2-MSA-1 and SARS-CoV-2-MSA-2 than random intervals of the same size (binomial test p-value < 1e-5). Table 1 lists these regions, which we term the **SARS-related-conserved** regions. These genome regions are conserved across the betacoronavirus sequences in SARSr-MSA-1 and have at least 99% sequence conservation across whole-genome sequences from the SARS-CoV-2 outbreak as of March 18, 2020 (SARS-CoV-2-MSA-2).

**Table 1:**
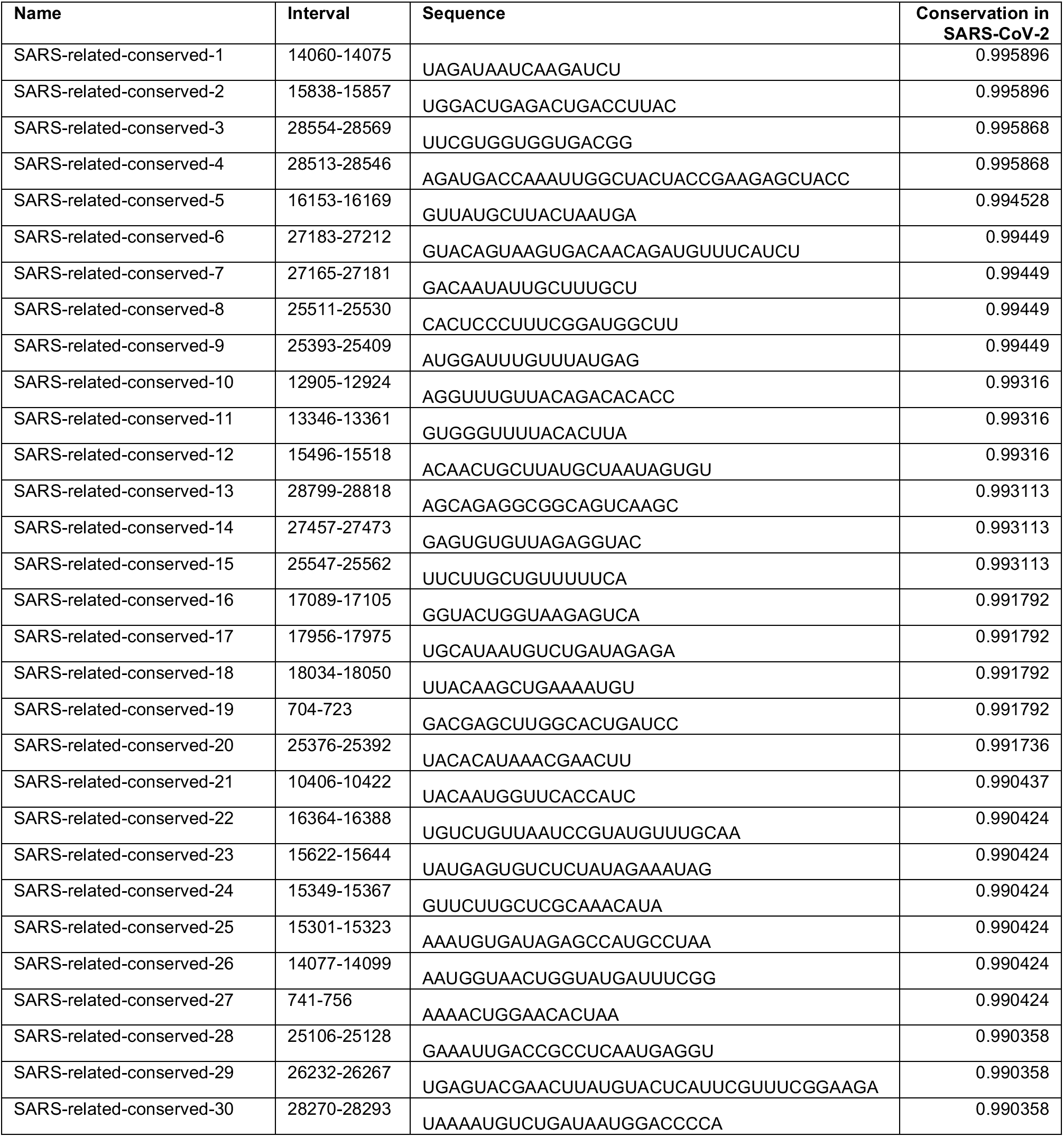
SARS-related-conserved. Conserved regions across SARSr-MSA-1 and SARS-CoV-2-MSA-2. All intervals are at least 90% conserved across the SARS and bat coronavirus sequences in SARSr-MSA-1, have length at least 15 nucleotides, and have every position at least 99% conserved in current GISAID SARS-CoV-2 sequences (SARS-CoV-2-MSA-2). Sequence intervals are relative to the reference genome NC_045512.2.

Conservation percentages for SARSr-MSA-1, SARSr-MSA-2, SARS-CoV-2-MSA-1, and SARS-CoV-2-MSA-2 are included in Supplementary File 1. We expect that some diagnostic and therapeutic strategies will benefit from focusing on conserved regions across a broad range of betacoronaviruses, whereas others may benefit from focusing on regions conserved only in SARS-CoV-2; we revisit the latter category of SARS-CoV-2-unique regions below.

### Predictions for structured regions in SARS-CoV-2

The intrinsic RNA structure of a conserved genome region is of interest in current medical research (Fig. 1B). On one hand, stable secondary structure domains are candidates for harboring stereotyped 3D RNA folds that present targets for small-molecule drug therapeutics. On the other hand, if an RNA region is sufficiently unstructured to allow binding by hybridization probes, antisense oligonucleotides may be used to disrupt these functional structures. Such unstructured stretches may also be more likely to be accessible to diagnostic and antiviral interventions including standard RT-PCR assays.

We used two approaches to make predictions for conserved structured regions in SARS-CoV-2. First, we predicted RNA structures centered on the most sequence-conserved regions of SARS-related betacoronavirus genomes (alignment SARSr-MSA-1). For each conserved stretch (at least 15 nucleotides long, 100% sequence conservation) along with 20 nucleotide flanking windows, we predicted maximum expected accuracy (MEA) secondary structures using Contrafold 2.0.^18^ We then sought to rank sequences based on the predicted probability that the RNA folds into the MEA structure and not other structures. For this ranking, we used the estimated Matthews correlation coefficient (MCC) from each construct’s base-pairing probability matrix.^19^ We note here that while MCC is often used in the RNA structure modeling literature to assess agreement of a prediction with a reference structure, we here use the metric to assess how tightly concentrated the ensemble of predicted secondary structures is to a single predicted secondary structure, the MEA structure. An MEA structure with a higher estimated MCC is expected to have unpaired and paired bases that better align with the construct’s predicted ensemble base-pairing probabilities, lending support to the single-structure MEA prediction. In Fig. 2 regions A-E, we display the five conserved regions with the top maximum expected accuracy (MEA) secondary structures as ranked by the estimated MCC (all regions listed in Supplementary File 1). Regions D and E occurred within the 5’ UTR and correspond to known SARS-related virus stem loops SL5a and SL2, respectively. Interestingly, region A is close to but does not overlap with the frameshifting stimulation element; it lies 200 nucleotides downstream of the FSE and could perhaps be involved in a more elaborate structure, as has been described for human coronavirus 229E and other coronaviruses.^20^

We also sought independent methods to identify thermodynamically stable and conserved RNA structures, without initially guiding the search to focus on extremely sequence-conserved genome regions. We made predictions for structured regions using RNAz^21^, beginning with the betacoronavirus alignment SARSr-MSA-1. RNAz predicts structured regions that are more thermodynamically stable than expected by comparison to random sequences of the same length and sequence composition (z-score), and additionally assesses regions by the support of compensatory and consistent mutations in the sequence alignment (SCI score). These two criteria are combined into a single P-score, which when tested empirically on a set of ncRNAs produced a false-positive rate of 4% at a P>0.5 cutoff and 1% at a P>0.9 cutoff. To predict structured regions across the full viral genome, we scanned the SARSr-MSA-1 alignment in windows of length 120 nucleotides sliding by 40 nucleotides, predicted all RNAz hits in the plus strand at a P>0.5 cutoff, clustered the resulting hits to generate maximally contiguous loci of the genome with predicted structure, and filtered results to only include loci with at least one window with a P>0.9 structure prediction.

The RNAz approach led to the prediction of 44 structured genome loci comprising 117 windows with predicted structure (P>0.9), with these loci covering 46% of the SARS-CoV-2 genome (Figure 3). We found that five canonical RNA structures (the frameshifting element, the 3’ UTR pseudoknot, the 3’ UTR hypervariable region, 5’ UTR SL2-3, and 5’ UTR SL5) were present in these loci. Additionally, conserved SARS-CoV-2 regions overlap significantly with predicted RNAz loci, with 62 of 78 SARS-CoV-2 conserved intervals at a 97% sequence cutoff overlapping by at least 15 nucleotides with RNAz loci. This enrichment is statistically significant (p-value<0.001 from comparisons to 10,000 random placements of conserved intervals). This enrichment also holds when considering overlaps with conserved regions from SARSr-MSA-1; 124 of the 229 SARSr-MSA-1 conserved intervals at a 90% conservation cutoff overlap by at least 15 nucleotides with RNAz loci (p-value 0.0038). This analysis potentially expands the set of conserved structural regions of SARS-CoV-2 beyond known Rfam families and those noted in the literature (full set of RNAz loci in Supplementary File 1). Top-scoring structured windows from RNAz that overlap with conserved sequence regions in SARS-CoV-2-MSA-2 for at least 15 nucleotides are included in Table 2; we termed these **SARS-CoV-2-conserved-structured** regions. Overlapping intervals between the RNAz predictions and conserved sequence regions in SARSr-MSA-1 are included in Supplementary File 1.

**Figure 3:**
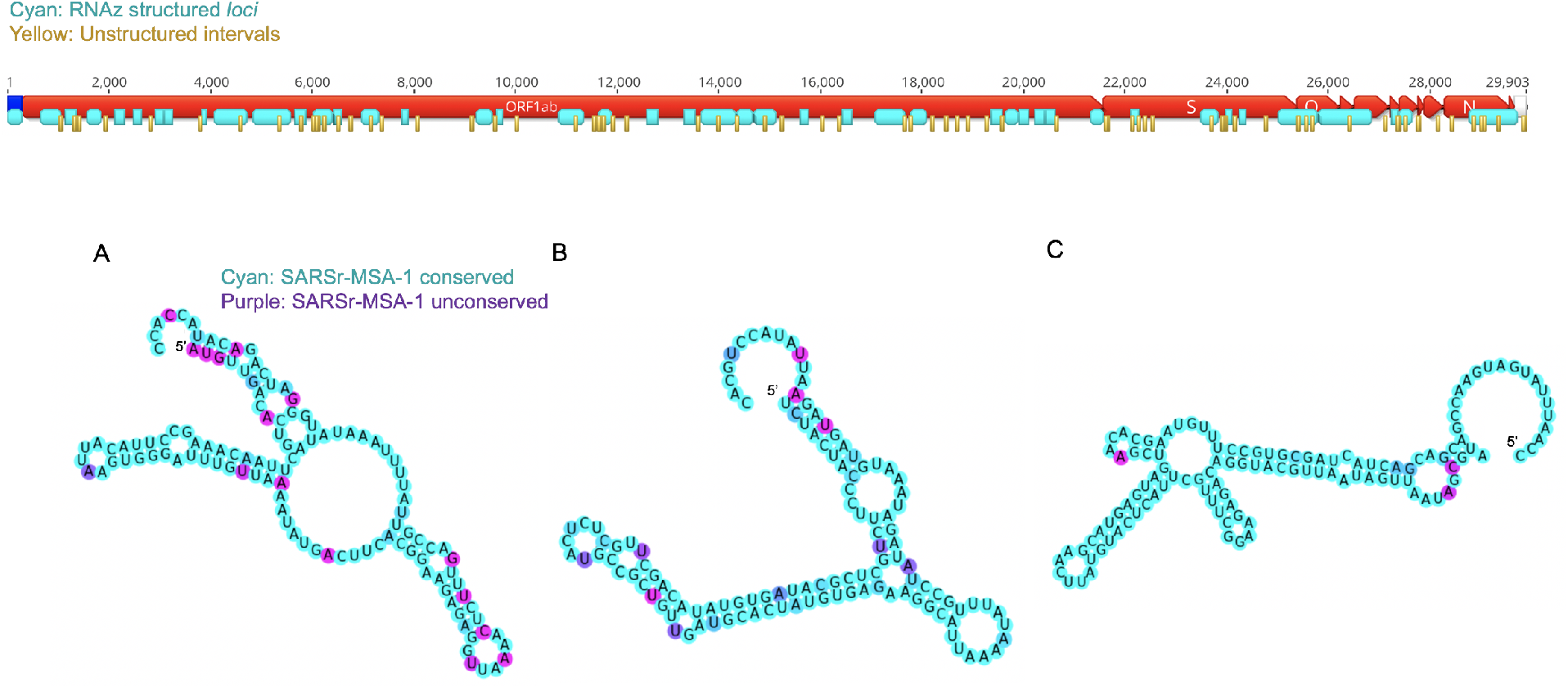
Structured (cyan) and unstructured (yellow) intervals on the genome ORFs for SARS-CoV-2, predicted from RNAz and a Contrafold 2.0 analysis, respectively. (A-C) highlight the three secondary structures for windows that do not overlap with known Rfam or literature-annotated structures with the highest P-value scores from RNAz (all P>0.9). These windows are located at genome positions 14207-14366 (A), 17126-17245 (B), and 26176-26295 (C). Secondary structures are colored by sequence conservation (cyan = more conserved, purple = less conserved). Figures prepared in Geneious^41^ and draw_rna (https://github.com/DasLab/draw_rna).

**Table 2:**
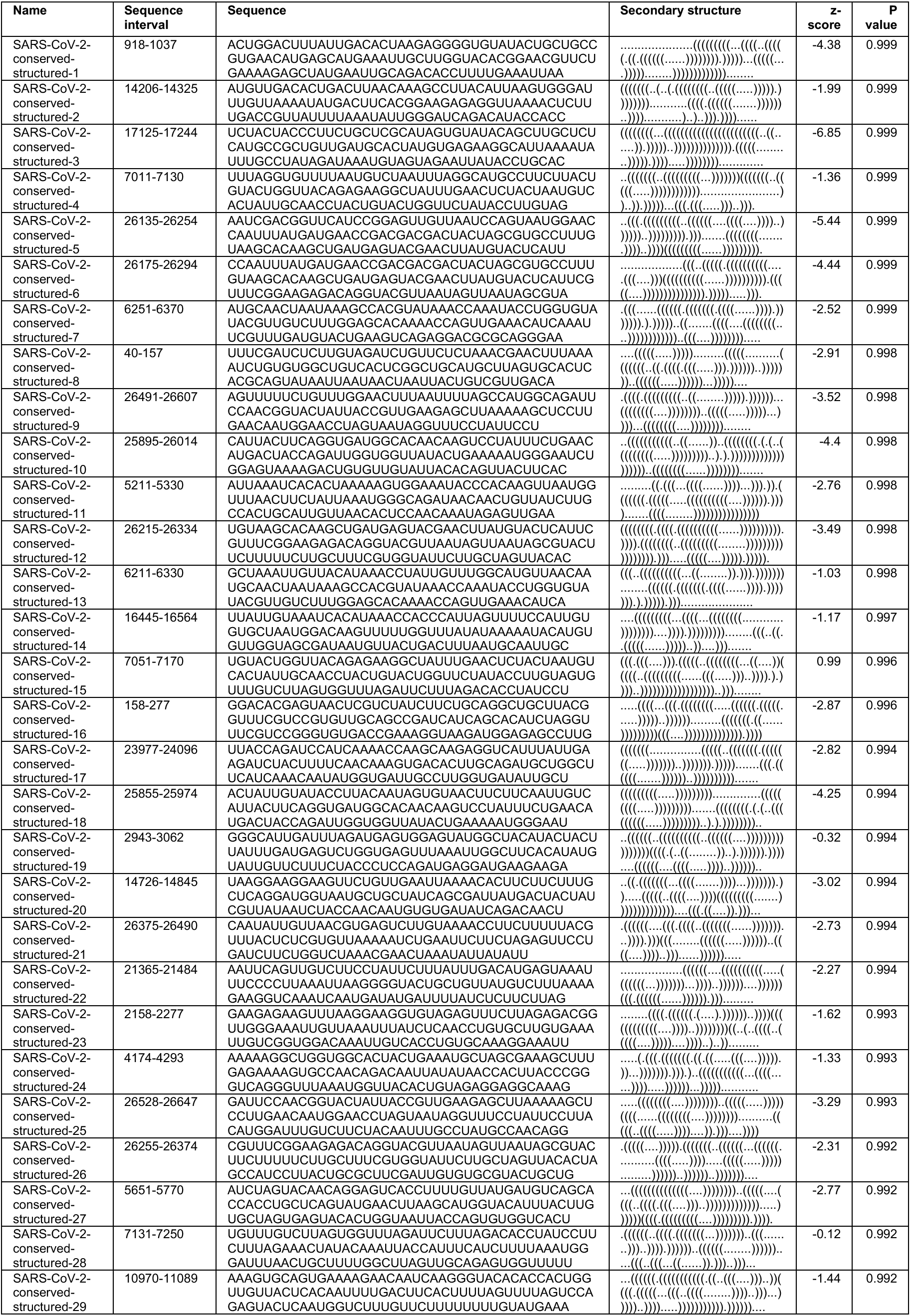
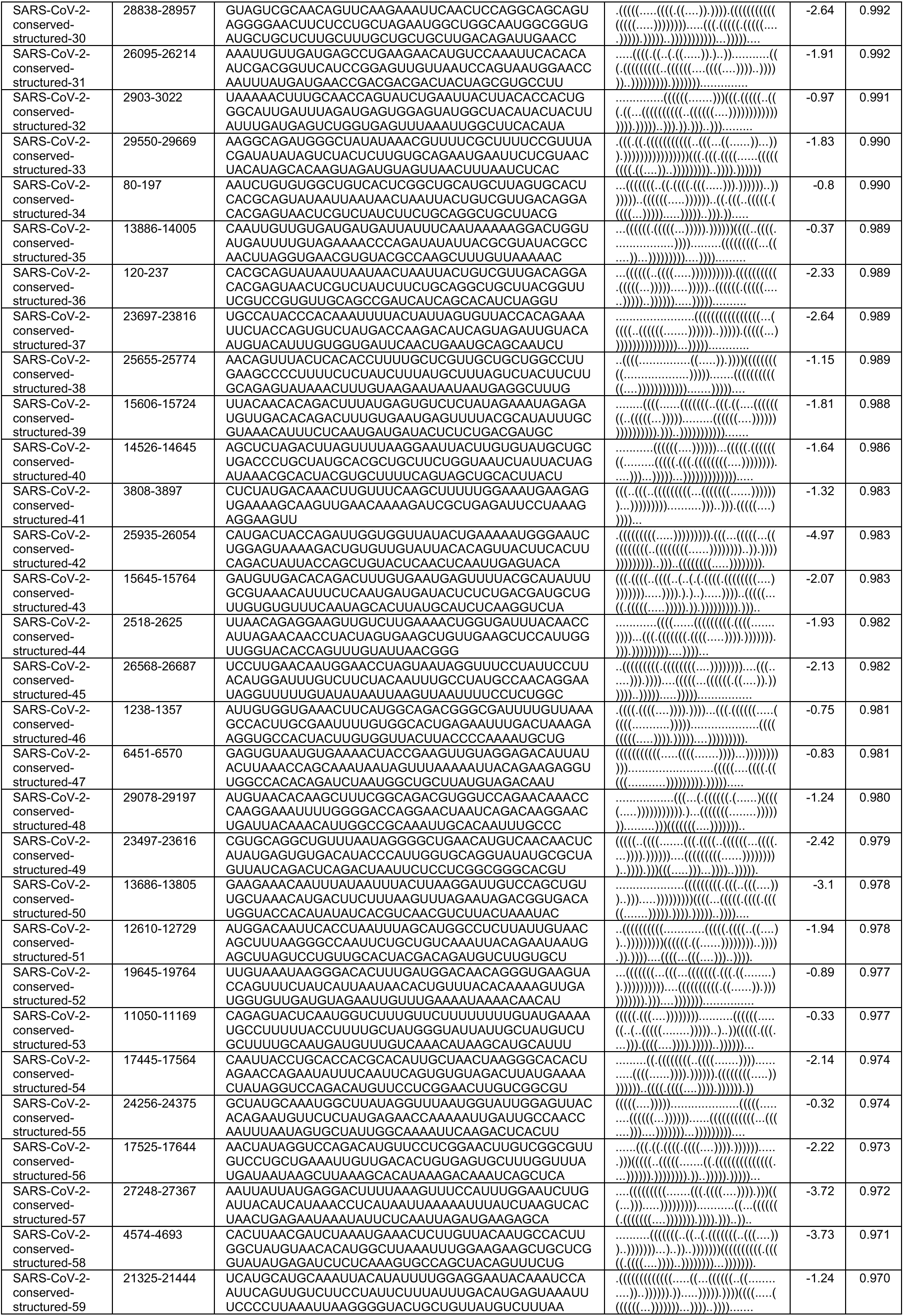
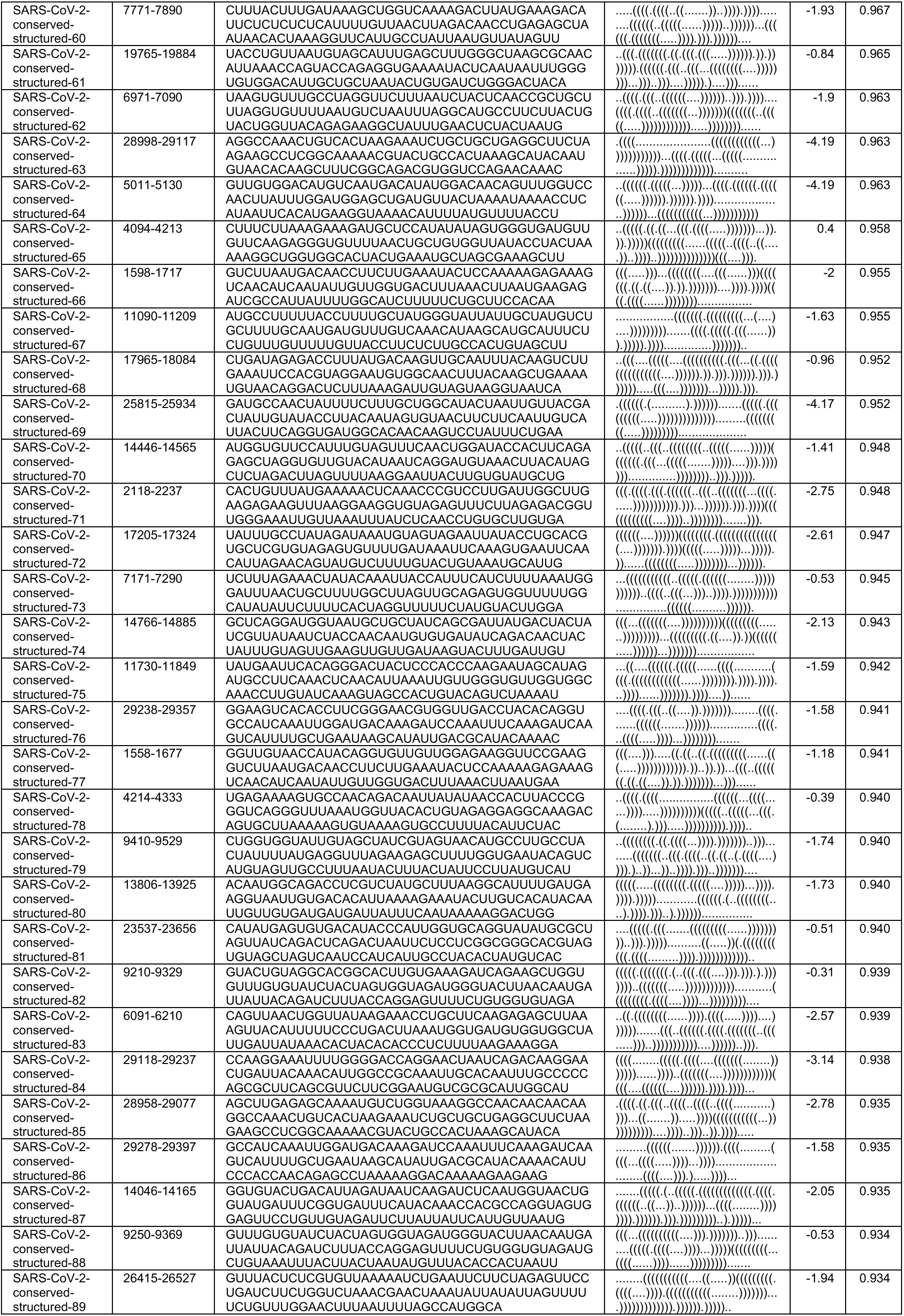
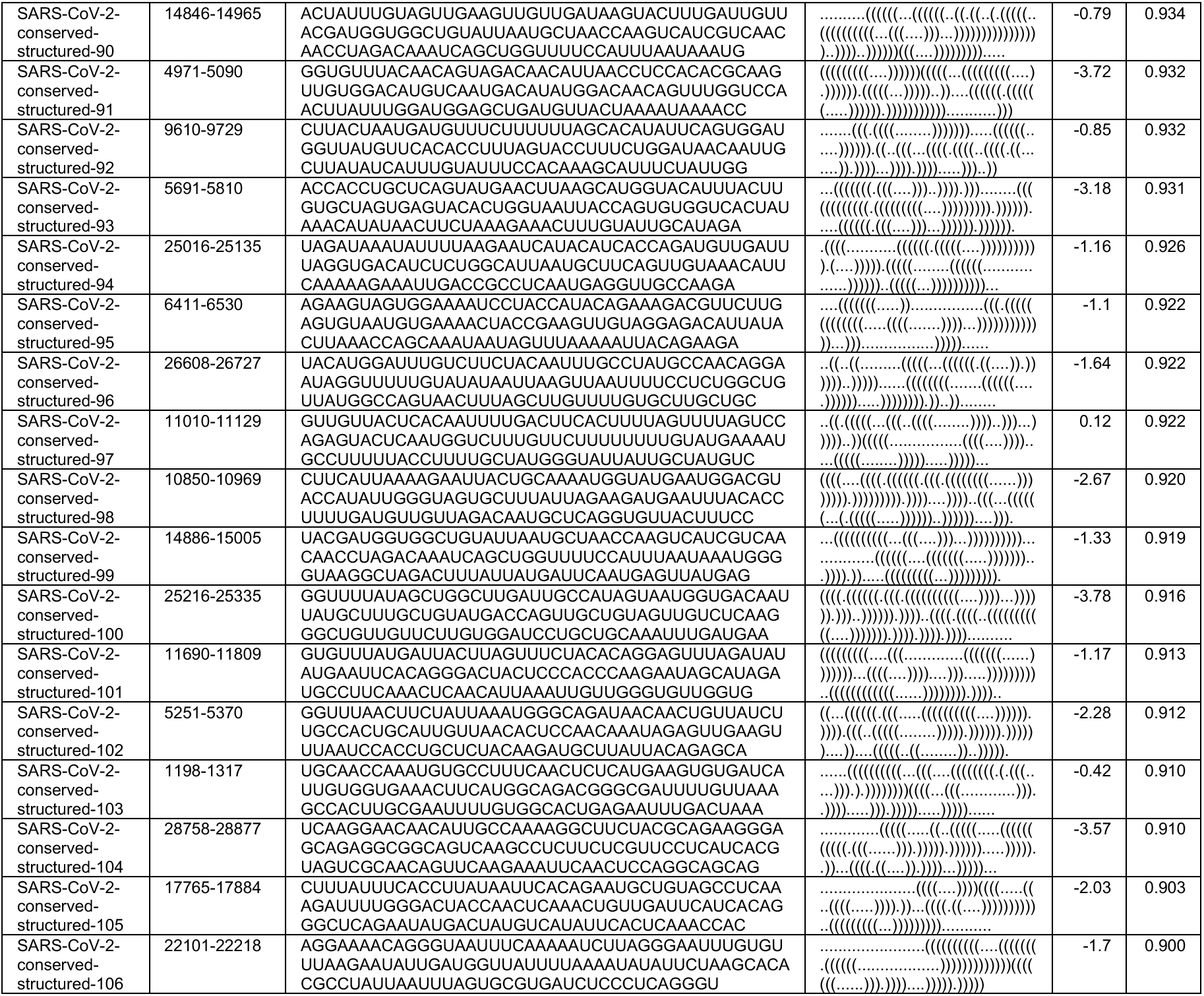
SARS-CoV-2-conserved-structured. RNAz windows as scored by the P-value (P>0.9) that overlap with conserved intervals from SARS-CoV-2-MSA-2 (97% conservation cutoff) by at least 15 nucleotides. Sequence intervals are relative to the reference genome NC_045512.2.

We sought to further check structured windows reported by RNAz using orthogonal approaches. First, we explored using R-scape to make structure predictions with covariation signal in the sequence alignments. However, we found that the SARSr-MSA-1 alignment had insufficient variation to detect conserved base pairs with covariation, lacking alignment power for all genomic windows.^22^ Second, we validated structured window predictions with alifoldz, a program that calculates a z-score for an alignment window by comparing the window’s consensus minimum free energy structure to that of random shuffled alignments. To mirror the RNAz analysis above, we scanned through windows of length 120 nucleotides sliding by 40 nucleotides. We chose a z-score cutoff of – 2.69, which kept only 1% of windows when running alifoldz on all shuffled windows across the genome. This approach led to predicting 228 alifoldz structured windows, overlapping with 104 of the 117 RNAz structured windows (P>0.9 cutoff). This overlap is statistically significant (p-value<1e-05). RNAz structured windows supported by alifoldz analysis are highlighted in Supplementary File 1.

### Conserved unstructured regions of SARS-CoV-2

We additionally located conserved regions of the viral genome predicted to *lack* structure, as such regions may be desired targets for some diagnostic and therapeutic approaches (Fig. 1C). We scanned the SARS-CoV-2 reference genome in windows of length 120 nucleotides sliding by 40 nucleotides, and for each window, we predicted the base-pair probability matrix with Contrafold 2.0, using these probabilities to assemble average single-nucleotide base pairing probabilities across the genome. In Figure 3, we display the 76 stretches of the genome of length at least 15 nucleotides where every base has average base-pairing probability at most 0.4.

It is interesting to note that some structured 120 nucleotide windows reported by RNAz include these unpaired stretches. A potential explanation for this observation is that such regions encode for well-defined, conserved RNA structures that themselves harbor long unpaired loops to recruit proteins, distal RNA elements, or other molecular machinery.

Overall, we find that 58 of these unpaired stretches have at least 15 nucleotides of overlap with sequence regions that are at least 97% conserved in SARS-CoV-2-MSA-2 (Fig. 5). These unpaired stretches termed **SARS-CoV-2-conserved-unstructured** regions are listed in Table 3 (overlaps with SARSr-MSA-1 are included in Supplementary File 1.)

**Table 3:**
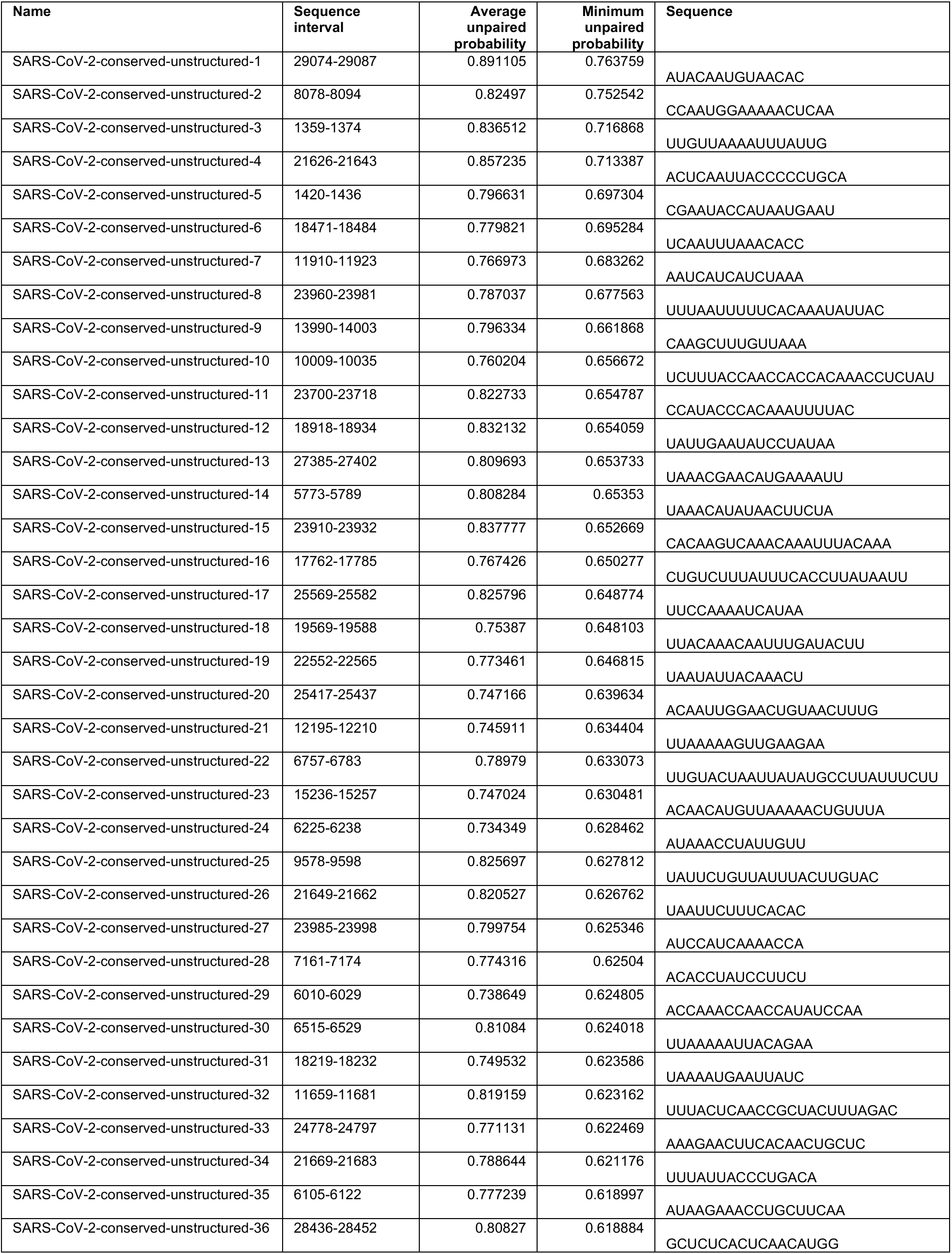
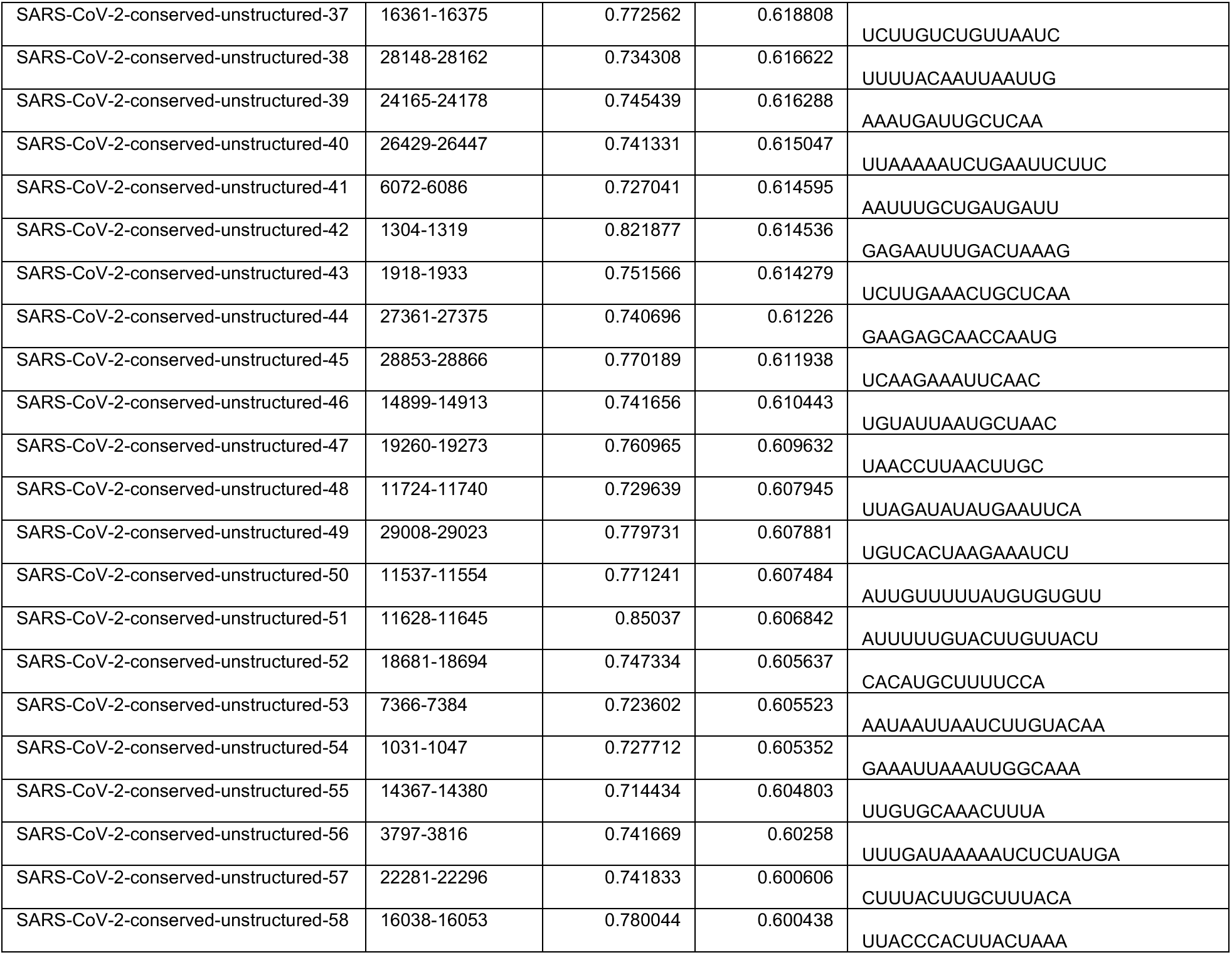
SARS-CoV-2-conserved-unstructured. Top unstructured regions (ranked by minimum unpaired probability over the interval, stretch of at least 15 nt) that overlap with conserved intervals from SARS-CoV-2 for at least 15 nt at a 97% sequence conservation cutoff. Sequence intervals are relative to the reference genome NC_045512.2.

As an orthogonal check for the unstructured intervals predicted using Contrafold 2.0 base-pairing probabilities, we used Vienna’s RNAplfold to compute unpaired probabilities for each genome position. In general, we found that RNAplfold predicted lower unpaired probabilities than Contrafold 2.0, with only 10 intervals of length at least 15 nucleotides having at least 0.6 probability of being unpaired, in contrast with the 76 stretches predicted by Contrafold 2.0. Nevertheless, we found that 9 of the 10 intervals predicted by Vienna’s RNAplfold overlap with unpaired intervals predicted from our Contrafold 2.0 analysis (regions listed in Supplementary File 1.)

### Secondary structure models for canonical structured regions of SARS-CoV-2

Currently known RNA structures that recur across betacoronaviruses provide potential starting points for therapeutic development targeting the SARS-CoV-2 RNA genome. Here, we include secondary structures for the 5’ UTR (Figure 4a), frame-shifting element (Figure 4b), and 3’ UTR (Figure 4c) for SARS-CoV-2, built by analyzing homology to literature-annotated structures in related betacoronaviruses. We additionally include computer-readable secondary structures in Table 4 and Supplementary File 1. A brief review of salient secondary structure features in these regions and their putative functional roles in the betacoronavirus life cycle follows.

**Figure 4.**
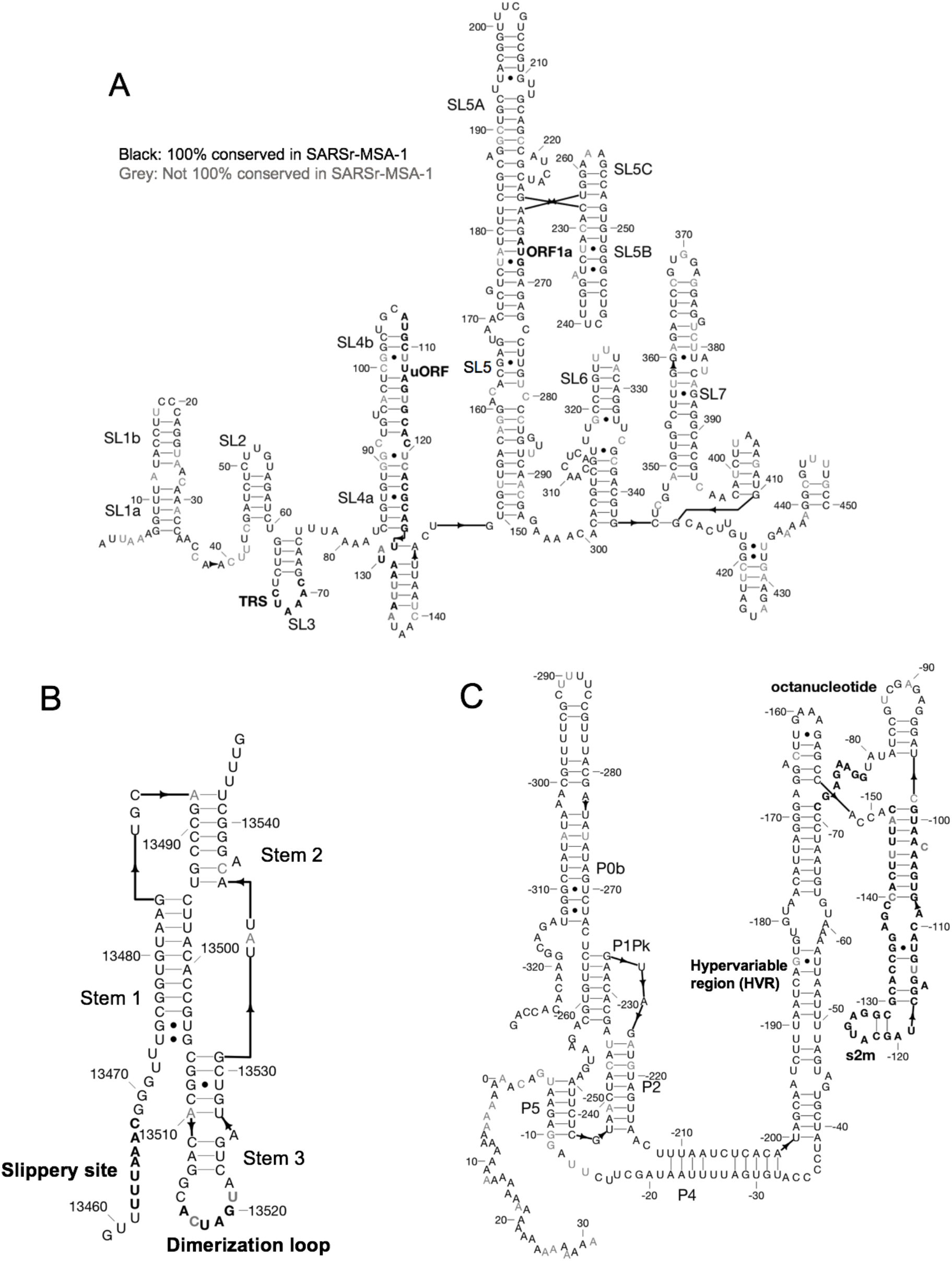
Secondary structure diagrams for A) 5’ UTR, B) Frameshift element, C) 3’ UTR. Nucleotides are black if 100% conserved in the SARS, bat, and SARS-CoV-2 sequences in SARSr-MSA-1, and grey otherwise. Special labeled domains are in boldface. Structures are based primarily on manual identification of homology with literature coronavirus structure models. Note that numbering in (C) is relative to 3’ end of virus sequence. Figures prepared in RiboDraw (https://github.com/ribokit/RiboDraw).

**Table 4:**
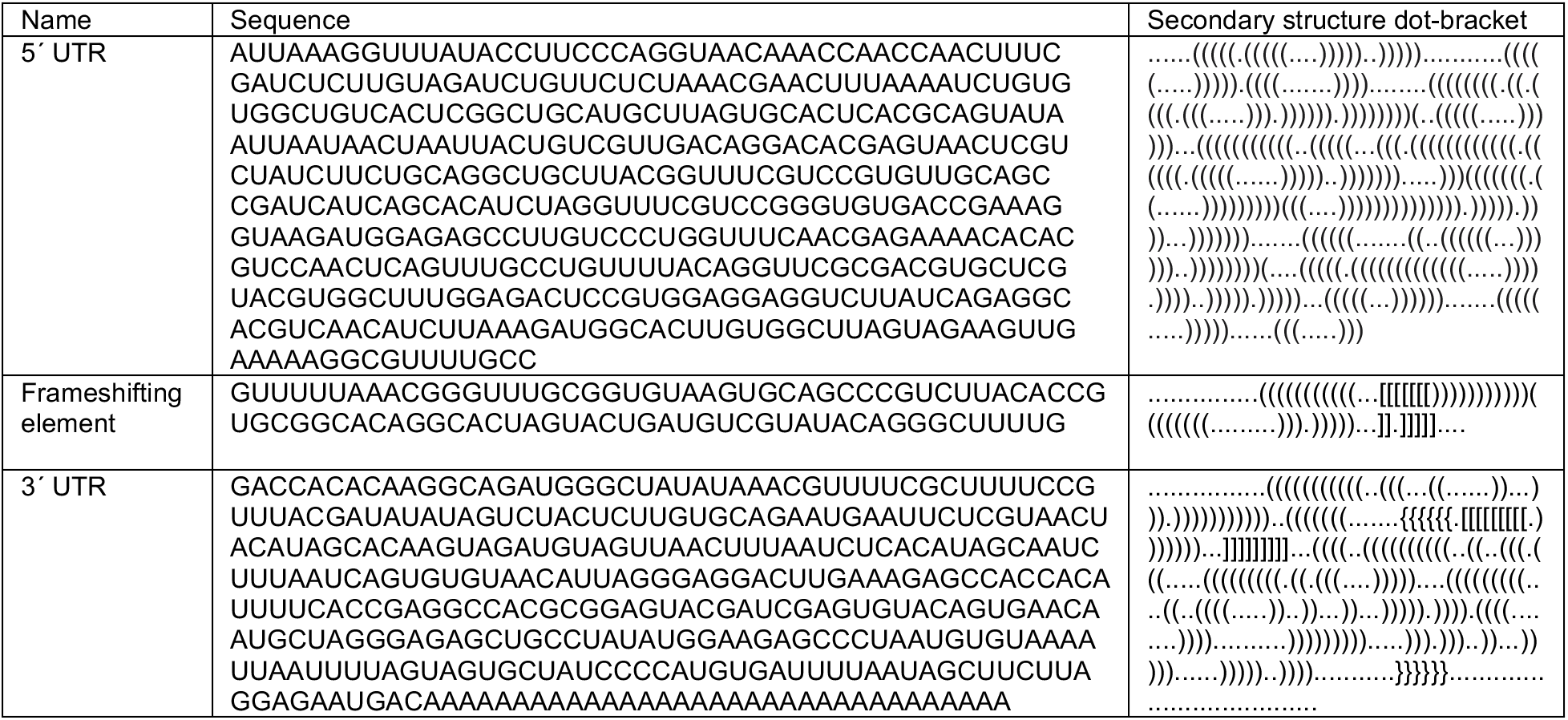
Sequences and secondary structure dot-brackets for key structured genome regions of SARS-CoV-2.

The 5’ UTR includes five confident stem-loop structures (SL1-SL5), with structures verified by chemical mapping experiments in related coronaviruses.^9, 16^ SL1 and SL2 are conserved across betacoronaviruses, with SL2 having the highest sequence conservation across the 5’ UTR.^13^ The high A-U base-pairing content in the SARS-CoV-2 SL1 sequence and the bulged nucleotides align with prior reports that SL1 is relatively thermodynamically unstable to allow for the formation of long-range interactions.^23^ SL2 has been shown to be critical for subgenomic RNA synthesis, with mutations in its conserved pentaloop retaining the production of genome-sized RNA, but not subgenomic RNA segments.^24^ SL3, conserved only in betacoronaviruses, presents the transcription-regulating sequence (TRS) that base pairs with one of several complementary sequences in nascent negative-sense strands in a ‘copy-choice mechanism’ that gives rise to discontinuous transcription of subgenomic mRNAs.^6^ SL4 contains a short upstream ORF, here labeled uORF, which precedes the first longer ORF1ab of the genome. The uORF leads to attenuated transcription of ORF1ab that appears helpful but is not essential for viral replication.^13^ SL5 has been implicated in a potential role in viral packaging, and contains the AUG start codon for long ORF1ab which encodes the viral replicase/transcriptase polyprotein. The SARS-CoV-2 SL5 domain has common features with the domain in other group IIb betacoronaviruses, for instance including UUCGU pentaloops on SL5a and SL5b, and a GNRA tetraloop on SL5c.^9^ Prior DMS-probing data for Stem 5 in SARS-CoV aligned with the proposed SL5a,b,c structures.^9^ Two additional stems (SL6 and SL7) are predicted from computer modeling here, but prior literature has not established whether such stems embedded in the coding region are functionally important across betacoronaviruses.

The frameshifting element (FSE) is located in ORF1ab and is involved in regulating a (−1) ribosomal frameshift event that is necessary for producing ORF1b. The FSE consists of a conserved pseudoknot structure that regulates the rate of ribosomal frameshifting at an upstream slippery site.^7^ This domain is nearly exactly conserved between SARS-CoV and SARS-CoV-2, suggesting a similar mechanism for ribosomal pausing and slippage between the two viruses.^25^

The 3’ UTR contains various domains critical for regulating viral RNA synthesis and potentially translation. The most 5’ region of the 3’ UTR includes a switch-like domain involving mutually exclusive formation of a pseudoknot and stem-loop, both of which are essential for viral replication with putative roles in establishing the kinetics of RNA synthesis.^6, 26^ The hyper-variable region (HVR) is not essential for viral RNA synthesis, as this can be removed while allowing for viral replication in tissue culture; however, viruses without this domain have lower pathogenicity in mice.^27^ This domain contains a completely conserved octonucleotide sequence with unconfirmed functional significance. The stem-loop II-like motif (s2m) is another subregion of the HVR that is conserved in SARS-CoV-2 and other coronaviruses. A crystal structure of the SARS s2m domain has been shown to be homologous to an rRNA loop that binds translation initiation proteins, leading this domain to have a proposed role in recruiting host translational machinery.^28^ The domain has been proposed to be a selfish element due to its recurrence in numerous virus families outside the *Coronaviridae,* but its function is not well understood.^29^

## Discussion

Understanding the RNA structure of the SARS-CoV-2 genome can guide RNA-targeting interventions and diagnostics. Here we have presented an initial analysis of RNA sequence conservation across betacoronaviruses and current SARS-CoV-2 sequences, predictions for structured and unstructured domains of the viral RNA genome, and homology-derived secondary structure models for classic structured elements of the SARS-CoV-2 genome: the 5’ UTR, the frame-shifting element, and the 3’ UTR. By filtering for sequences that have more than one of these properties, we have curated three sets of RNA genomic regions of potential interest for further structural analysis, which we have termed the SARS-related-conserved, SARS-CoV-2-conserved-structured, and SARS-CoV-2-conserved-unstructured sets. Fig. 5 gives a more extensive presentation of how these sets overlap. Our hope is that these steps will provide useful starting points for efforts to develop antivirals and diagnostics that depend on targeting either structured or unstructured viral genomic regions.

**Figure 5.**
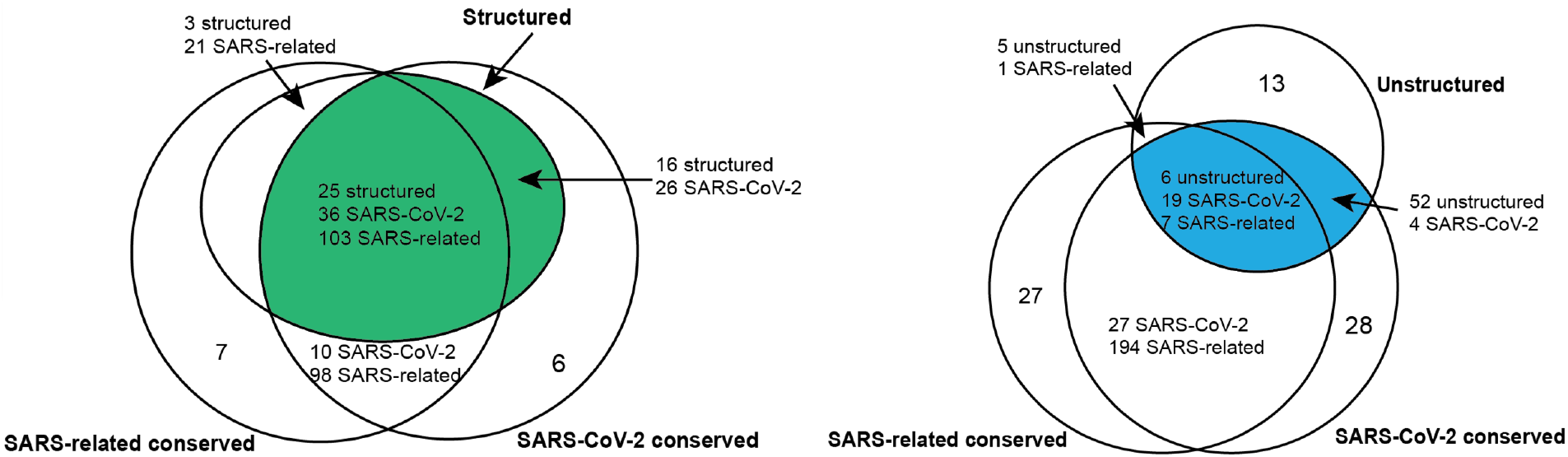
We depict the predicted number of structured, unstructured and conserved intervals for a choice of sequence conservation cutoffs. The SARS-related conserved intervals are all regions of at least 15 nucleotides with each position at least 90% conserved across an alignment of SARS, bat coronavirus, and SARS-CoV-2 sequences (SARSr-MSA-1). The SARS-CoV-2 intervals are regions of at least 15 nucleotides with each position at least 97% conserved across an alignment of currently available SARS-CoV-2 sequences (SARS-CoV-2-MSA-2). Structured intervals are *loci* predicted from RNAz with some *loci* containing multiple RNAz windows, and unstructured intervals are stretches of at least 15 nucleotides where all bases have base-pairing probability at most 0.4. All interval intersections are required to have at least 15 nucleotide overlaps, with the number of overlapping intervals listed for each interval type involved in the intersection. Top-scoring structured intervals conserved in SARS-CoV-2 sequences (green) are listed in Table 2. Top-scoring unstructured intervals conserved in SARS-CoV-2 sequences (blue) are listed in Table 3.

The abundant RNA structures involved in the replication cycle of betacoronaviruses present ample opportunities for therapeutic development, but our analysis is not complete. First, while homology to prior structures annotated in betacoronaviruses lends some confidence to the 5’ UTR, frame-shifting element, and 3’ UTR secondary structures presented here, there may still be some inaccuracies. For instance, SL6 and SL7 in the 5’ UTR are built based on computer modeling. As another example, the frame-shifting element structure presented here differs in two base pairs compared to that presented by Kelly and Dinman.^25^ Additional biochemical and genetic verification, particularly through compensatory mutagenesis, will be needed to further assess these structures.

Beyond the secondary structures highlighted here, prior work has pinpointed a variety of RNA-RNA interactions important to the betacoronavirus life cycle. Long-range interactions between the 5’ UTR and 3’ UTR have been implicated in RNA synthesis, for instance with mutations in the 5’ UTR SL1 only supporting viral replication when co-evolving with specific mutations in the 3’ UTR.^13, 30^ Such long-range RNA-RNA interactions would be missed in our analyses above which have focused on shorter windows. In related coronaviruses, RNA structures that act as packaging signals have been identified in genomic ORFs.^13^ Such packaging signals may reside in regions identified here from our RNAz computational analysis or may have been missed.

We believe a more thorough structure/function analysis of the virus should be obtainable with recent experimental technologies that integrate multidimensional chemical mapping, electron microscopy, and computer modeling.^31–32^ New candidates for structured RNA elements in SARS-CoV-2 could play various functional roles, perhaps regulating viral packaging, replication, RNA synthesis, or translation initiation. To further improve these structure predictions, information about protein-binding events should be integrated, and biochemical assays can be conducted with these proteins present or even in cells. Accounting for protein-binding events will be critical to completing a picture of accessible and structured RNA sites.

The secondary structures predicted here present a reasonable starting point for 3D modeling of RNA-only structures in various regions, including the 5’ UTR stem loop 5, the frame-shifting element, the 3’ UTR pseudoknot, and the 3’ stem-loop II-like motif. Furthermore, these regions and novel predicted structures can serve as candidates for RNA-only structure determination. Such 3D structures have the potential to reveal well-defined 3D folds with conserved binding domains for small-molecule drugs, potentially presenting alternative approaches for targeting SARS-CoV-2.

## Methods

### Conservation analysis

Three alignments for SARS-related viruses were prepared:

1. SARSr-MSA-1 was generated by re-aligning the sequences curated by Ceraolo and Giorgi^10^ with MUSCLE^33^ using default alignment settings, excluding all non-reference genome copies of the SARS-CoV-2 sequence and excluding MERS sequences JX869059.2 and KT368829.1.
2. SARSr-MSA-2 was generated by downloading the MSA provided by BLAST for the top 100 complete genome sequences closest to the SARS-CoV-2 reference genome.
3. SARSr-MSA-3 was generated by obtaining all complete betacoronavirus genome sequences available from the NCBI database, removing mutually similar sequences using a 99% cutoff with CD-HIT-EST^34^, and computing an MSA with Clustal Omega^35^ using default settings.

Two alignments for SARS-CoV-2 sequences were prepared:

1. SARS-CoV-2-MSA-1 was generated by downloading the MSA provided by NCBI for the 103 whole-genome SARS-CoV-2 sequences deposited as of 03-18-20.
2. SARS-CoV-2-MSA-2 was generated from 739 GISAID^17^ sequences, including all sequences described in the Nextstrain project metadata file as of 03-18-20 (https://github.com/nextstrain/ncov). Sequences were aligned using MAFFT^36^ with the --add flag to add GISAID sequences to seed alignment SARS-CoV-2-MSA-1, and the sequences in SARS-CoV-2-MSA-1 were subsequently removed to avoid duplicates.

All alignments are included in the associated GitHub repository: https://github.com/DasLab/SARSCoV2_Secstruct_Cons.

### Analysis of structured elements

To identify regions in the SARS-CoV-2 reference genome NC_0405512.2^11^ that were matches to Rfam^14^ families, we used Infernal^15^ to build covariance models from Rfam families RR0164, RR0165, and RR0507 with cmbuild, and we ran cmscan to find hits with an E<1e-04 threshold.

MEA structures were computed using Contrafold 2.0^18^ for 20 nucleotide flanking windows around conserved intervals in SARS-CoV-2. To estimate the Matthews correlation coefficient of these single-structure predictions, we computed the pseudo-expected Matthews correlation coefficient as described in Hamada, et al.^19^ Base-pairing probability matrices were computed Contrafold 2.0, and these were then used to calculated the expected number of true positive, true negative, false positive, and false negative base pairs. These computations were carried out using the Arnie package (https://github.com/DasLab/arnie).

RNAz^21^ structures were predicted in windows of the SARS-CoV-2 genome using the SARSr-MSA-1 alignment. We used rnazWindow.pl to compile alignment windows across SARSr-MSA-1 with at least 4 sequences in each window, using a window size of 120 nucleotides sliding by 40 nucleotides, and using default settings otherwise. RNAz hits were computed at the P>0.5 threshold for the forward strand with z-scores computed without a shuffled sequence background for efficiency, using the --no-shuffle flag. The resulting RNAz structured windows were then clustered with rnazCluster.pl, filtered with rnazFilter.pl at a P>0.9 threshold, and sorted with rnazSort.pl.

We ran alifoldz^37^ on the same genome windows used with RNAz above, again using SARSr-MSA-1. The alifoldz z-score computations were calculated for the forward strand only with alifoldz.pl. We additionally calculated alifoldz z-scores for alignment windows that were shuffled with shuffle-aln.pl to assess background z-scores, determining that 1% of shuffled alignment z-scores were less than −2.69.

We computed alignment powers with R-scape^38^ to assess the potential for using SARSr-MSA-1 for covariation analysis. We generated Stockholm alignment files with biopython^39^ for windows of 120 nucleotides each sliding by 40 nucleotides. For each window, we ran R-scape with the –fold flag to predict new structures, obtaining estimates for the power of each base pair in the predicted structure (here, power is the expected sensitivity for detecting base pairs given the number of substitutions in the alignment at that base pair). We then averaged across base pairs in each structure to obtain the alignment power as described in Rivas, et al.,^22^ noting that all windows’ alignment powers fell below the 0.10 threshold used by Rivas, et al.^22^ to distinguish low-power from high-power alignments.

### Analysis of unstructured elements

To obtain probabilities that each genome position in SARS-CoV-2 was unpaired, we computed basepairing probability matrices with Contrafold 2.0^18^ in windows of 120 nucleotides sliding by 40 nucleotides, and for each genome position, we summed the probabilities of pairing with all potential partners. We then averaged these nucleotide pairing probabilities across all windows that nucleotide was present in. Additionally, RNAplfold^40^ was run with window size 120 nucleotides, producing another set of unpaired probabilities for each position in the genome.

### Code and data availability

Code used for the conservation, structured, and unstructured analyses above can be found at the GitHub repository: https://github.com/DasLab/SARSCoV2_Secstruct_Cons. The repository additionally includes alignment files, Rfam families and covariance models, and output from the RNAz, R-scape, alifoldz and RNAplfold analyses.

## Acknowledgments

The authors acknowledge support from the National Science Foundation Graduate Research Fellowship Program under grant no. 1650114 (R.R.), a Stanford Graduate Fellowship (I.N.Z.), and NIH grant MIRA R35 GM122579 (to R.D.). We would like to thank Dr. Rachel H. Saluti, Dr. Edward A. Pham, and Dr. Jeffrey S. Glenn for providing advice on the conserved, structured intervals desirable for antisense oligonucleotide design; Hannah Wayment-Steele for useful discussions on secondary structure modeling; Andrew M. Watkins for updates to the RiboDraw software for rendering secondary structures; and Dr. Paul Gardner for providing rapid reviews of the initial preprint of this manuscript on bioRXiv.

